# Photosynthesis in the biomass model species *Lemna minor* displays plant-conserved and species-specific features

**DOI:** 10.1101/2023.04.25.538322

**Authors:** Monique Liebers, Elisabeth Hommel, Björn Grübler, Jakob Danehl, Sascha Offermann, Thomas Pfannschmidt

## Abstract

*Lemnaceae* are aquatic freshwater plants with extraordinary high growth rates. We have studied selected physiological and molecular photosynthesis properties of the duckweed *Lemna minor* and compared these to the terrestrial model species *Arabidopsis thaliana. Lemna* and *Arabidopsis* plants grown under identical light intensities displayed similar photosynthesis characteristics, however, *Lemna* exhibited slighty better quenching efficiencies pointing to improved light utilization in the duckweed. Western-immuno-blot analyses of representative photosynthesis proteins suggest various post-translational modifications in *Lemna* that might be associated to this. Phospho-threonine phosphorylation patterns of thylakoid membranes uncovered differences between the two species. Testing the photosystem II antenna association of *Lemna minor* in dark and light by 77K chlorophyll fluorescence emission experiments, however, revealed a typical association as reported in terrestrial plants. High light stress experiments causing photoinhibition and subsequent recovery from it were not substantially different in *Lemna* when compared to *Arabidopsis*. We hypothesize that the molecular differences in *Lemna* photosynthesis proteins are associated with evolutionary adaptations to the aquatic life style and ultimately with the high growth rates. We also developed a video imaging approach for *Lemna* multiplication at high magnification that will be useful to assess the impact of external factors on *Lemna* photosynthesis and growth.

## Introduction

The duckweed *Lemna minor* is a member of the *Lemnaceae*, a small family of cosmopolitan aquatic plants found in freshwater ecosystems consisting of five genera with in total 36 species [1]. Despite their simple two-dimensional morphology comprising a small number of leaves laying on the water surface and a species-specific number of roots, they are true vascular plants belonging to the group of monocotyledons. *Lemnaceae* usually multiply by vegetative propagation without sexual recombination although they are able to flower and to sexually reproduce under specific conditions. They typically grow in calm sweet water ponds or lakes and, under ideal conditions, can exhibit extraordinary growth rates with biomass doubling times of less than 48 h rendering them the fastest growing plants on Earth [2]. Because of these particular properties *Lemnaceae* are highly interesting for a plethora of biotechnology applications including for instance the production of food for cattle and human, production of biofuels as well as a high variety of biomolecules such as vitamins, carotenoids, starch (and many others), or as bioassay organisms in toxicity tests and also in waste water management [3-5].

Because of its high biotechnology potential, the research interest in *Lemnaceae* has largely increased in the last 10 years. Most potential applications are directly or indirectly connected to the extraordinary growth properties of these plants. However, the reasons why *Lemnaceae* exhibit these high growth rates remain largely not understood until now, especially at the molecular level. Potential reasons discussed include their special ecological niche with unlimited water supply, the high availability of nutrients dissolved in the water column or the reduced limitation in light harvesting through competition for light with neighbouring plants or through self-shading. The primary energy source for growth in all plants is photosynthesis. The high growth potential of *Lemnaceae* thus could be associated to specific molecular properties, e.g. in photosynthesis, not yet identified. The *Lemnaceae* have been used for many ground-breaking studies in plant biochemistry and physiology in the 1950-1990ies including photosynthesis research [6-8] but were largely neglected afterwards with the rise of genetically more easily tractable model systems such as *Arabidopsis thaliana* or *Chlamydomonas reinhardtii* [3]. Many aspects of our understanding of photosynthesis in *Lemnaceae*, thus, are either technically out-of-date or have never been investigated in depth. In addition, the recent advances in genome sequence information provides potential novel information not available in the early studies on photosynthesis of *Lemnaceae* [9].

A recent study demonstrated that *Lemna gibba* displays a high plasticity in its response to changing growth light intensities indicating that this duckweed can easily adapt to highly varying light intensities [10]. There are recent reports indicating that also *Lemna minor* can grow effectively over a range of light conditions [11,12]. Here, we have studied a number of important general and selected molecular photosynthetic properties of *Lemna minor* by a combination of physiological and biochemical approaches. We report commonalities as well as specific differences to the terrestrial model organism *Arabidopsis* which may provide first hints on evolutionary molecular adaptations associated to the aquatic lifestyle and high growth potential of *Lemnaceae*.

## Results

### Imaging of *Lemna minor* growth

We have grown *Lemna minor* under sterile conditions on liquid medium as described [13]. Since sugars are known to induce inhibitory effects on photosynthesis gene expression in plants [14] we strictly avoided the addition of any carbon source in our growth media in order to assess the true photosynthesis potential of *Lemna minor*. To this end *Lemna minor* fronds were pre-cultured and maintained in the exponential growth phase by regular transfer between bottles of growth media (Fig. 1A). Fronds from these cultures were then used for further analyses. For physiological photosynthesis measurements *Lemna* fronds have been transferred to beakers or Petri dishes with identical growth medium, but in non-sterile conditions that allowed the removal of the lid for imaging of chlorophyll (Chl) fluorescence from top. *Lemna* fronds swimming on liquid medium, however, always showed some floating caused by micro-convections in air and medium that occurred even in a growth chamber. This floating often resulted in disturbances of the detected Chl fluorescence signals interfering with a trustful Chl fluorescence analysis. Prevention of floating could be achieved by filling the measurement vessel completely with *Lemna* fronds in such a way that the plants have no further space for any movement (Fig. 1B). However, *Lemnaceae* are known to be touch-sensitive and stop growing once a certain density is reached. Since this may have a feedback effect on photosynthesis we tested other ways to prevent the floating. For short-term imaging (in the range of min to few hours) we prevented any movement by pinching the roots of single fronds with a layer of staples on the bottom of a Petri dish. For long-term imaging (many hours to days) that may be disturbed in addition by growth-related movements we found the use of self-made plastid grids very efficient. It allowed growth of roots also of new fronds but prevented very effectively any growth-induced lateral movements and was suitable to record qualitative long-term high magnification videos of *Lemna* fronds during growth (Fig. 1C, D and Video S1). This set-up was found to exhibit a very high positional stability of single fronds allowing even the observation of stalk formation between mother and daughter fronds (Fig. 1D, white arrows). It furthermore allowed the direct determination of growth rates by detecting the recorded leaf area using ImageJ software (Fig. 1E) revealing growth rates comparable to those reported on the base of frond number [2].

**Fig. 1:**
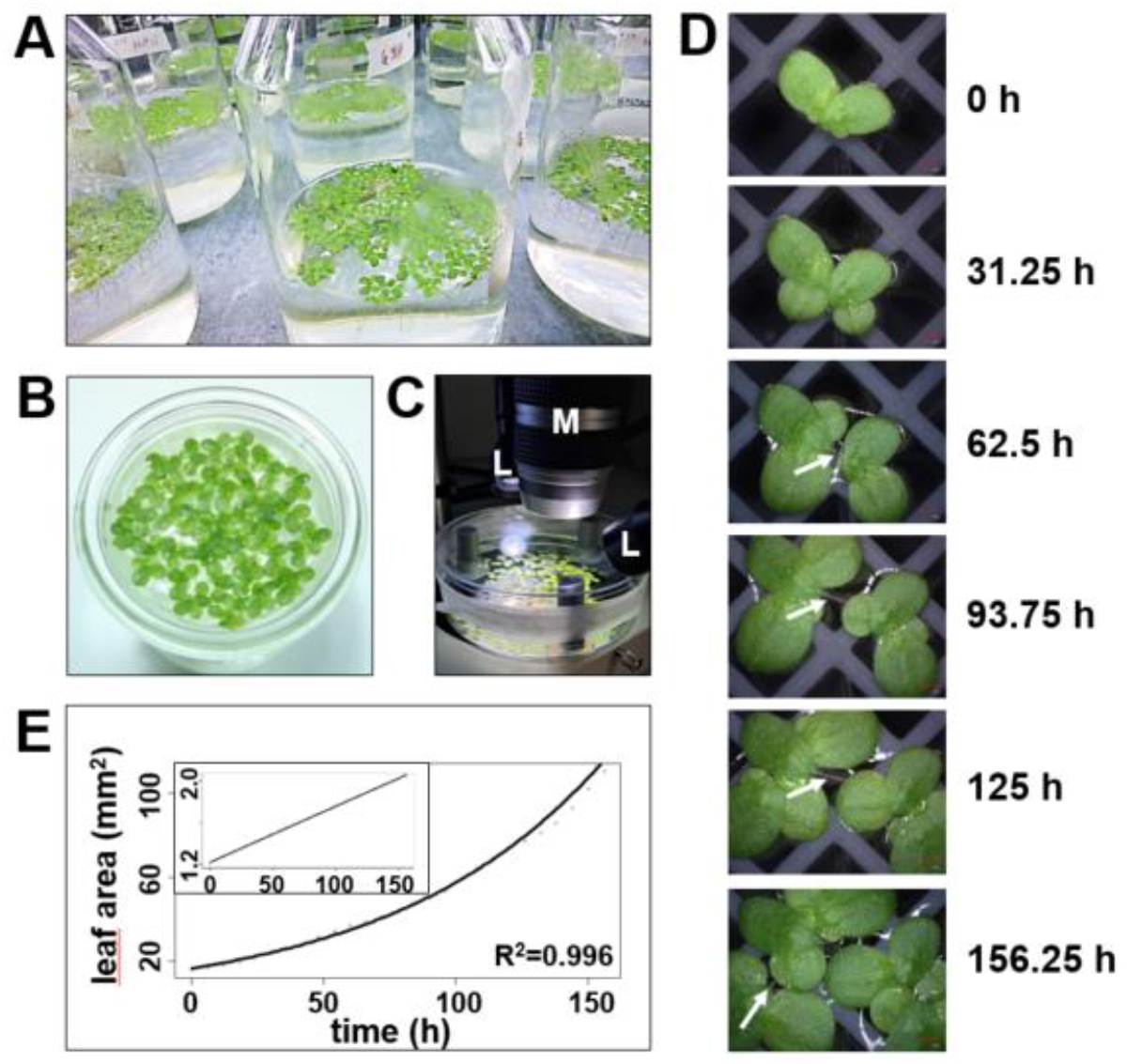
Imaging of *Lemna minor* growth characteristics in sugar-free liquid medium. **A)** Sterile mass pre-culture of *Lemna minor* cultivated in flasks with standard growth medium **B)** *Lemna minor* fronds from exponential growth phase placed in vessel for detection of Chi fluorescence from top. **C)** Set-up for time lapse movies recording *Lemna minor* growth. M: Objective of the digital microscope; L: Cold-white light source **D)** Time series of growth of *Lemna minor* fronds taken by a digital microscope with automatic z-axis zoom and software-based image construction. Red scale bar in bottom right comer of images represents 1 mm. White arrows indicate the stalk connecting mother and daughter fronds. The complete time lapse video of this series is available in the supplement. **E)** Growth curves of *Lemna minor* in sugar-free liquid medium determined as determined by the increase in leaf area detected in the time lapse video. The growth follows an exponential function (function of exponential regression: f(x)=16.477e^00125x^). The inset in the upper left corner displays a semi-logarhythmic representation.

### Basic photosynthesis characteristics of *Lemna minor*

We measured photosynthesis of single *Lemna minor* fronds with a pulse amplitude modulation (PAM) fluorometer following standard protocols [15,16] using the saturating pulse mode over a time period of 10 min (Fig. 2A). As reference we used *Arabidopsis thaliana* plants grown under identical conditions providing a comparison to a well characterized terrestrial model organism of photosynthesis (Fig. 2B). Dark-adapted *Lemna* fronds displayed a strong Kautsky effect after illumination appearing highly comparable to equally measured *Arabidopsis* plants. After the initial peak upon illumination, Chl fluorescence (F_t_) in *Lemna* declined rapidly and with the same velocity as in *Arabidopsis* indicating that the onset of photochemical as well as non-photochemical quenching processes followed the same kinetics as far as visible at the measured time scales. However, the steady state Chl fluorescence parameter F_s_ [17] in relation to F_m_ was slightly lower in *Lemna* than in *Arabidopsis* (0.06 in *Lemna* versus 0.21 in *Arabidopsis*) suggesting a more efficient transfer of electrons from PSII into subsequent processes (either photochemistry or dissipation). Also the intermediate rise in F_t_ typically resulting from the delayed onset of the Calvin cycle enzyme activities (reported for many vascular plants to occur) could be not observed with *Lemna* under these conditions. This may be indicative of a highly efficient or strong electron sink capacity of *Lemna minor* photosynthesis. In sum, the aquatic duckweed *Lemna minor* exhibited highly efficient quenching kinetics comparable or even superior to the well characterized terrestrial species *Arabidopsis thaliana* in these standardized photosynthesis measurements.

**Fig. 2:**
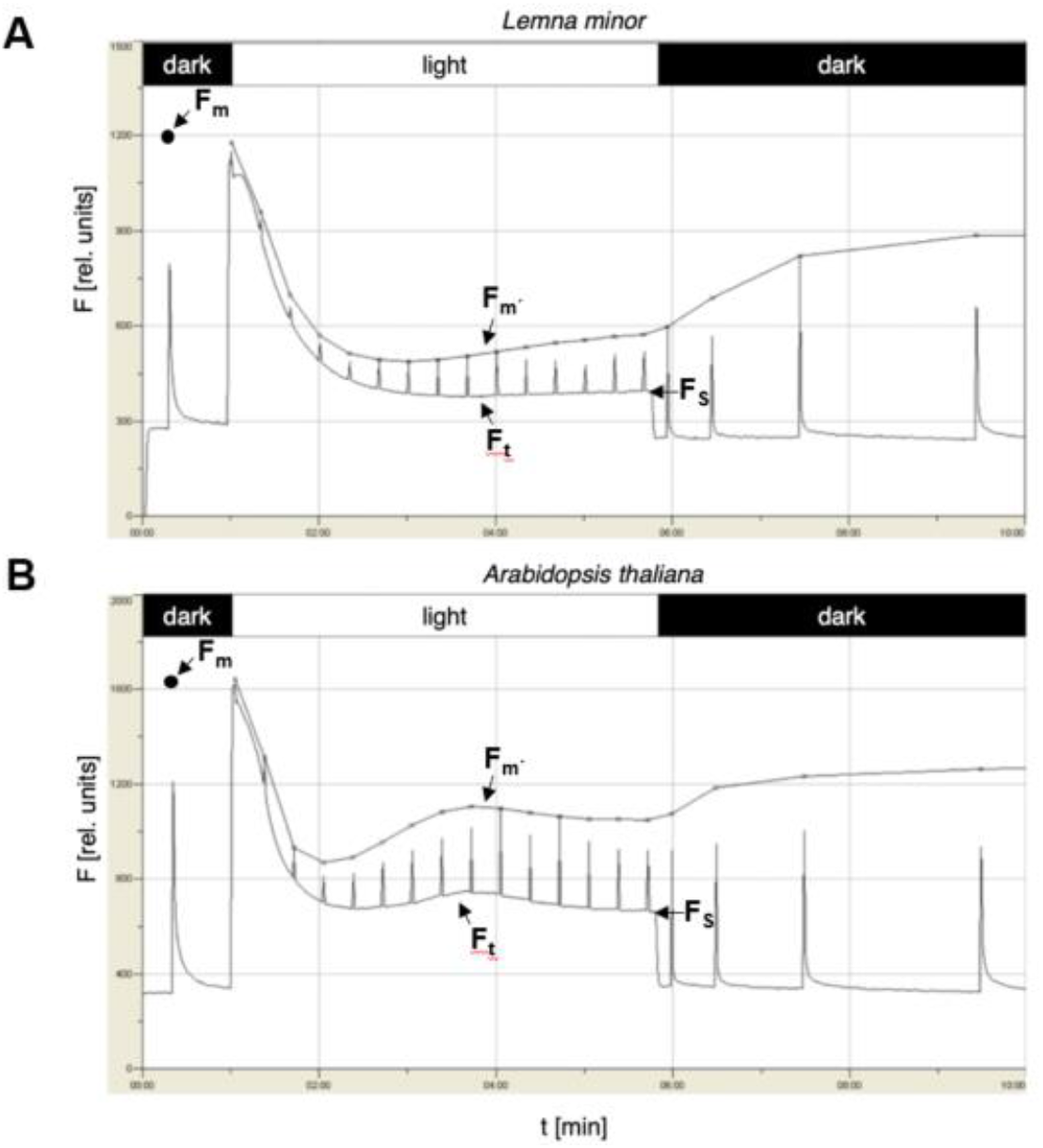
Photosynthetic performance of aquatic *Lemna minor* compared to terrestrial *Arabidopsis thaliana* as determined by Chi fluorescence. Chi fluorescence traces in a standard quenching analysis of dark-adapted plants upon illumination performed with a Junior-PAM device are shown. Fluorescence (F) is given in arbitrary units (rel. units). Light and dark phases during the measurements are indicated on top. corresponding time scale is given on bottom. **A)** Chi fluorescence from *Lemna minor*. Three independent biological replicates were done, one representative curve is given. For comparison the same type of measurement with identical technical settings performed with *Arabidopsis thaliana* **(B)** is given underneath. Note that the graphics of the measurement programme do not depict the saturation light pulses correctly. The true resulting maximal fluorescence is indicated by a line labelled with F_m_. F_m_: Maximal fluorescence; F_t_: Fluorescence in dependency of time t; Fs: Steady state fluorescence.

### Analysis of thylakoid proteins and their phosphorylation state

To get a deeper insight into the photosynthesis apparatus of *Lemna minor* in comparison to *Arabidopsis thaliana* we performed an initial structural characterization of photosynthesis thylakoid proteins by western-immuno-analyses using commercially available antisera. To this end we isolated total protein extracts from *Lemna* and *Arabidopsis*, separated them by SDS-PAGE and stained them by Coomassie (Fig. 3A). The general protein pattern of the two species was of similar appearance, but we also observed many variations in size and accumulation of individual proteins which represent either different species-specific proteins or modifications of homologous proteins. The protein-specific western immune analysis (Fig. 3B) revealed that all photosynthesis proteins in *Lemna* can be readily detected with antisera directed against *Arabidopsis* proteins.

**Fig. 3:**
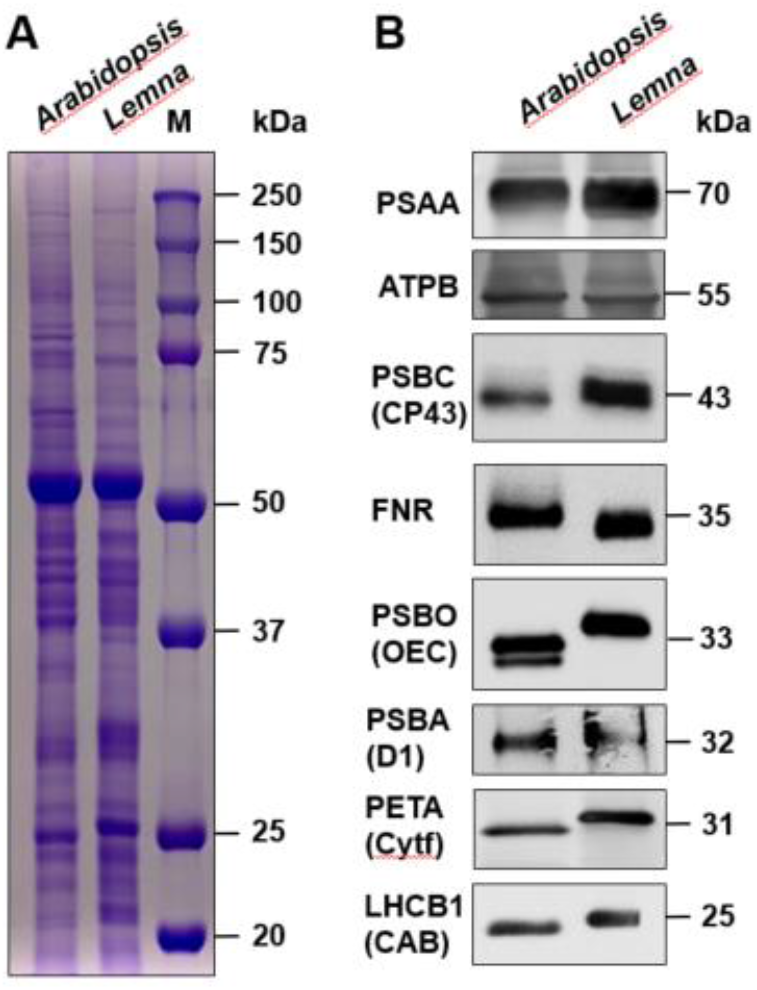
Comparison of total and photosynthesis proteins from *Arabidopsis thaliana* and *Lemna minor*. **A)** SDS gel (12%) electrophoresis of total protein extracts isolated from white-light-grown plants (identities indicated on top) stained with Coomassie. Marker (M) sizes are given in kDa at nght panel. **B)** Western-immuno-blot analyses of selected photosynthesis proteins. Protein identities are indicated in the left margin using both gene based and common (in parentheses) abbreviations, apparent molecular weight is given in the right margin. Loading of gel lanes was done as shown in A. Equality of loading was checked by Ponceau S staining of the membrane before immuno detection.

The general accumulation of these proteins was highly comparable between the two species suggesting that the general stoichiometries of the protein complexes within the photosynthetic electron transport chain do not differ between *Lemna* and *Arabidopsis*. However, we observed a number of specific differences regarding the migration behavior of those proteins. While PSAA (the apoprotein A of PSI), ATPB (the beta subunit of the ATPase) and PSBA (the reaction center protein D1 of PSII) did not display visual migration differences, PSBC (the PSII core protein CP43), PSBO (the 33kD protein of the oxygen evolving complex (OEC)), PETA (the cytochrome f subunit (Cytf) of the cytochrome *b*_*6*_*f* complex) and LHCB1, (one major chlorophyll A/B binding protein (CAB1) of the light harvesting complex of PSII (LHCII)) all displayed a larger apparent molecular weight in *Lemna*. Only FNR (ferredoxin-NADP+-oxidoreductase) revealed a smaller one. In addition, the signal from *Lemna* PSBC appears to consist of two very closely running proteins. We checked available genomic information about the proteins displaying differential migration behavior to elucidate whether these differences could be caused by evolutionary sequence variations in the corresponding genes (Text File S1). We found, however, very high degrees of conservation between these *Lemna* and *Arabidopsis* photosynthesis proteins indicating that the differences in migration behavior of the corresponding proteins cannot be simply attributed to sequence differences. This highly suggests the involvement of post-translational modifications of the respective proteins.

One major, if not the most important, post-translational modification of photosynthesis proteins is the phosphorylation at threonine residues in PSII proteins. Phosphorylation is further known to be a major cause for variation in migration behavior of proteins separated by SDS-PAGE. In order to test whether variations in the phosphorylation status might cause the observed differences in the migration behavior we examined the respective phosphorylation state of thylakoid membrane proteins by anti-phospho-threonine immune blotting. This technique is well established for terrestrial model plants but to our knowledge has never been employed in *Lemnaceae*. Our experiment was performed with material isolated from the light phase, a condition in which a high phosphorylation state of thylakoid membrane proteins should be expectable, especially in the growth light intensity we used. We observed indeed strong phosphorylation signals with both the *Arabidopsis* and the *Lemna* samples (Fig. 4). However, we identified also clear differences in the phosphorylation patterns (indicated by arrows in Fig. 4) that point to some specific differences in the phosphorylation states of both core as well as antenna proteins of PSII. These results support the idea that variations in post-translational modifications might be responsible for the observed differences in protein migration.

**Fig. 4:**
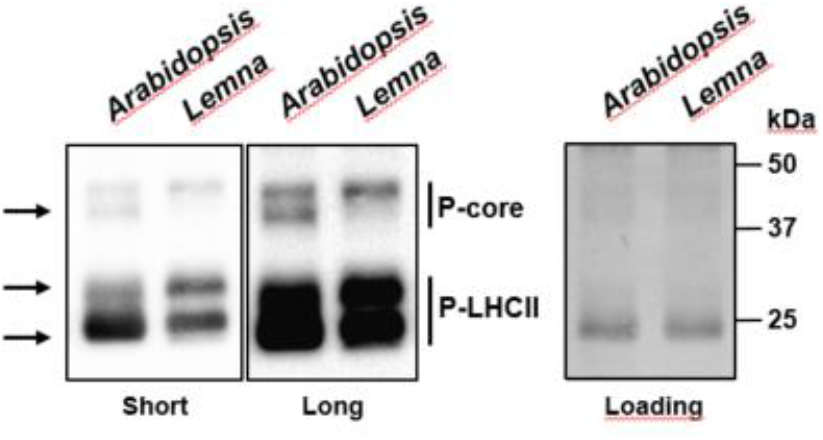
Phosphorylation state of thylakoid membrane proteins of *Lemna minor* in comparison to *Arabidopsis thaliana*. Protein amounts corresponding to 1 pg chlorophyll were separated by SDS PAGE and transferred to a nitrocellulose membrane *via* western blot. The membranes were incubated with anti-phospho-threonine antibodies to detect phosphorylated thylakoid membrane proteins using enhanced chemiluminescence (ECL). Signal detection was done for 7 sec (left panel, short) or 1 min (middle panel, long). Equal protein loading was tested by amido black staining of the membrane after ECL detection (loading, right panel). Sizes of marker proteins are given in the right margin in kDa Phosphorylated LHCII and PSI I core proteins are indicated in the right margin, differences in the phosphorylation signals by arrows at the left margin.

**Fig. 5:**
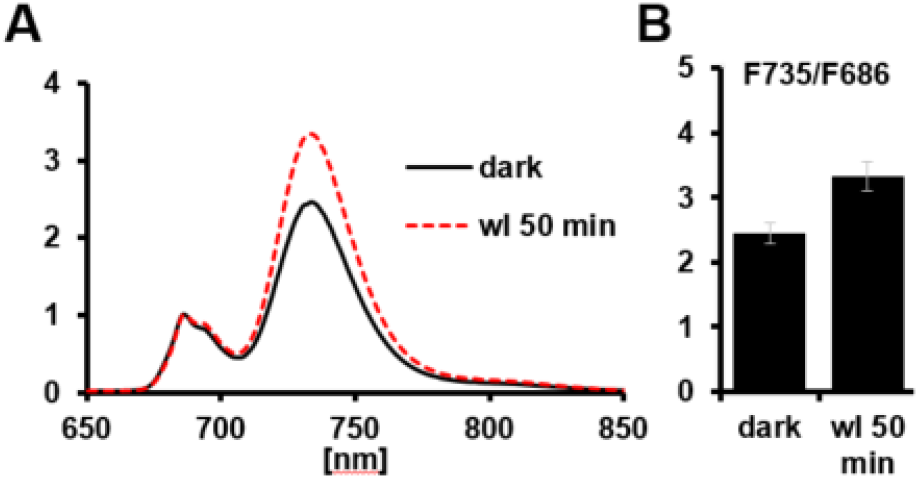
77K fluorescence spectra of *Lemna minor*. *Lemna minor* fronds were analysed for Chi fluorescence emission in the range of 650 - 850 nm excitation in liquid nitrogen. Measured plant materials were harvested at the end of the dark phase of the growth light regime or 50 min after the onset of white light illumination. All spectra were normalized to the fluorescence emission peak at 686 nm. **A)** Representative spectra for both conditions. **B)** Mean ratio of the fluorescence emission peaks at 735 and 686 nm (F735/F686) obtained from three independent replicates. SD is given.

### Photosynthesis antenna movements and photoinhibition in *Lemna minor*

Phosphorylation of thylakoid membrane proteins is functionally connected to a number of physiological responses that adapt the photosynthetic apparatus to varying environmental illumination conditions. Phosphorylation of LHCB proteins in terrestrial plants is responsible for the dynamic and reversible association of LHCII to either PSII or PSI that optimizes light harvesting (a process called state transitions) [18], while phosphorylation of PSII core proteins are known to be involved in the PSII repair cycle after high-light induced photoinhibition [19]. In order to assess the functionality of these processes in *Lemna minor* we conducted physiological experiments that are known to trigger corresponding responses in *Arabidopsis* [20,21]. Phosphorylation of the LHCII in *Arabidopsis* is catalyzed by the redox-sensitive thylakoid kinase STN7 that becomes activated in light upon a reduced plastoquinone pool. Our light intensity conditions chosen for growth are ideal to activate this kinase. We thus performed 77K Chl fluorescence emission experiments with *Lemna minor* fronds harvested from the dark phase (a condition with inactive kinase) and fronds harvested 50 min after the onset of white light (a condition expected to fully activate the kinase). In the dark sample we observed a ratio for the Chl fluorescence emission at 735 nm (originating from PSI) to the emission at 686 nm (originating from PSII) of around 2.4. In the sample collected 50 min after onset of light the ratio raised to around 3.2 indicating that the relative antenna size of PSI largely increased upon white light illumination. This light-induced increase in the F735/F686 ratio corresponds well to data reported for *Arabidopsis* [22] strongly suggesting that the *Lemna* homologue of *Arabidopsis* STN7 is working in a highly similar manner.

In order to assess photoinhibition and the recovery from it in *Lemna minor* we performed a high light treatment of isolated fronds placed into wells of a micro-titer plate each filled with liquid growth medium. To maximize the comparability of results *Arabidopsis* plants in growth pots were placed alongside the *Lemna* fronds and in an identical distance to the light source. We used a high intensity LED array that provided 1800 μmol photons m^-2^ s^-1^ white light at the height of leaf surface to induce photo-damage of PSII. Light-induced photoinhibition typically becomes apparent by a decrease in the F_v_/F_m_ value since the D1 protein of PSII becomes damaged and less PSII reaction centers are available for photochemistry. The return of F_v_/F_m_ to normal levels then indicates the repair of the damaged PSII reaction centers. We recorded this Chl fluorescence parameter with a 2D imaging PAM in which both species can be assessed at the same time (Fig. 6). Both *Lemna* and *Arabidopsis* displayed a comparable progressive decline of the F_v_/F_m_ value during the course of the high light treatment (Fig. 6A; 0h, 0.25h, 0.5h, 1h, 2h). Upon return to low intensity growth light conditions both plant species recovered to full extent and with comparable kinetics (Fig. 6). In sum our analyses indicate that the observed differences in the phosphorylation patterns of thylakoid membrane proteins of *Lemna* and *Arabidopsis* do not result in quantitative differences in physiological acclimation of photosynthesis to varying illumination, at least under the conditions we have tested.

**Fig. 6:**
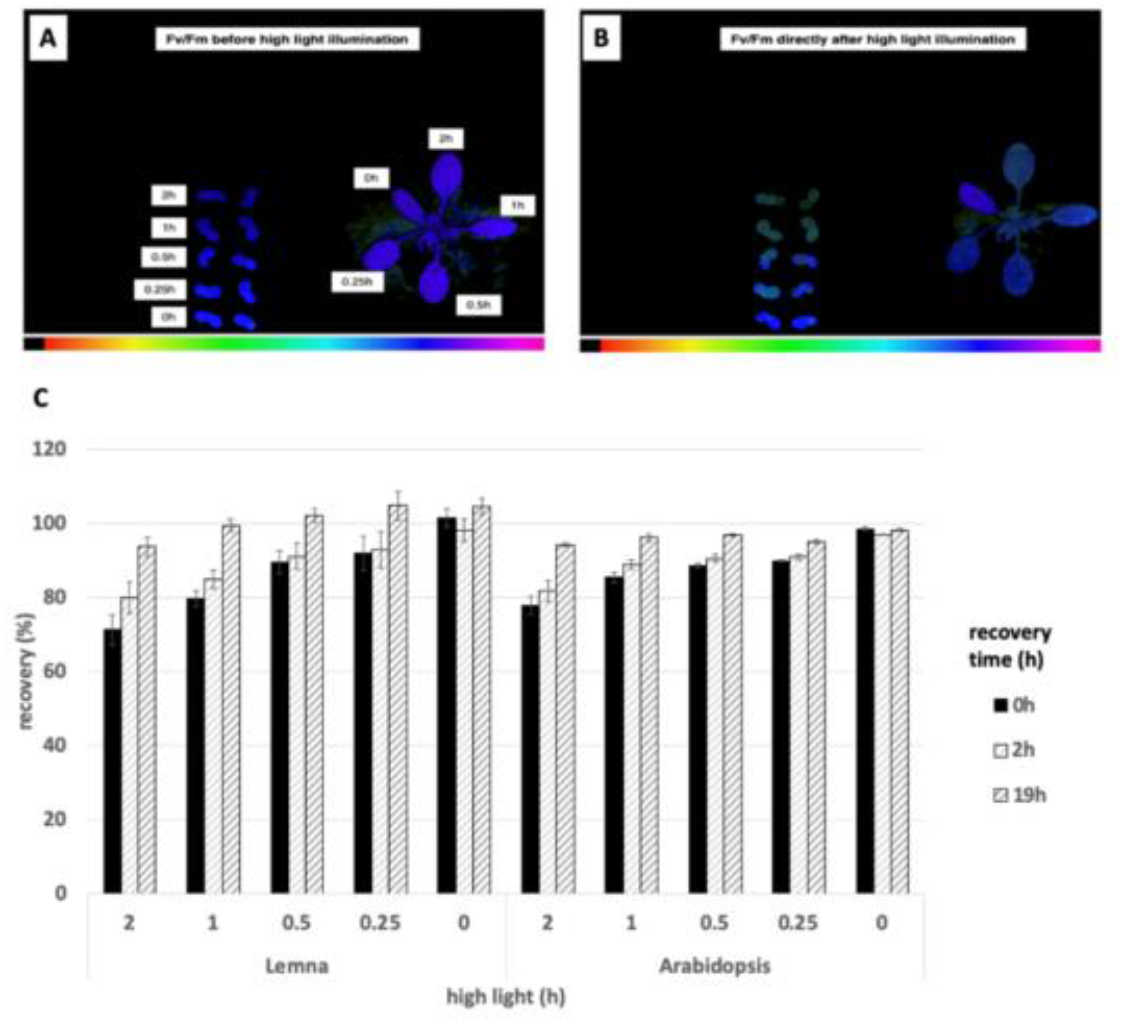
High-light induced photoinhibition and recovery in *Lemna minor* and *Arabidopsis thaliana*. *Lemna minor* (*L.m.*) and Arabidopsis thaliana (*A.t.*) were exposed to high light (HL) (1800 pmol nr^2^ PPFD) and tested for photoinhibition and recovery from it. **A, B)** Representative images of the F_v_/F_m_ values recorded before (A. reference) and directly after **(B)** high light (HL) treatment. Duration of illumination for the HL treatment is indicated by the numbers in **A**. F_v_/F_m_ values are color coded from black indicating F_v_/F_m_ = 0 to purple indicating F_v_/F_m_ = 1 (see color gradient bar below the images). **C)** Duration of HL treatment is indicated on x-axis. F_v_/F_m_ was measured before the HL treatment (as reference) and immediately (Oh. black bars). 2h (light grey bars) and 19h (dark grey bars) after return to growth light intensity. Recovery (rec. %) is expressed relative to the reference F_v_/F_m_ measured on a individual leaf/frond base. Results are averages of three biological independent replicates. Small black vertical bars indicate the standard error of the mean (SEM).

## Discussion

The ecological niche of the aquatic *Lemnaceae* is highly different from those of terrestrial plants with respect to water and nutrient availability, light competition, herbivore attack and many more stressors. Despite these differences the photosynthetic properties of *Lemna minor* and *Arabidopsis thaliana* appeared to be comparable at the physiological level in our study. We observed, however, small potential functional differences in our Chl fluorescence analyses that appear negligible at first sight, but may have a strong long-term impact on ecological adaptation and subsequent species survival. Future studies that include a wider range of physiological conditions, thus, will provide more detailed data allowing for building a more sophisticated functional model of *Lemna minor* photosynthesis. The presented study provides already a number of highly interesting aspects on which such future studies can build on.

Our study reported here on *Lemna minor* provided a first analysis of some properties known to be essential corner stones of molecular photosynthesis regulation in terrestrial plants. Phosphorylation of threonine residues in thylakoid proteins by redox-controlled kinases is a well-characterized and described regulation mode that controls a number of photosynthesis responses to the environment [23]. Despite the differences in thylakoid membrane phosphorylation (Fig. 4) *Lemna minor* displayed a well conserved light-dependent association pattern of LHCII to the photosystems (Fig. 5) and a kinetically comparable recovery response after photoinhibition with respect to the *Arabidopsis* control (Fig. 6). The critical phosphorylation residues for these regulatory processes thus are likely conserved. The specific differences at the molecular level that were detected between the two species (Figs. 3, 4) most likely represent evolutionary adaptations to the different ecological niches and may explain why photosynthesis in *Lemna* minor works efficiently at a water surface. The detected differences in SDS-PAGE migration behavior most likely are caused by specific post-translational modifications since our bioinformatics analyses revealed that sequence variations of corresponding genes are very minor and do not contain the potential to account for the observed differences. Our study was designed as a pilot project covering a number of basic photosynthesis parameters in order to elucidate whether more detailed analyses are worthwhile. The molecular differences we observed here are highly promising for our understanding of adaptations in plant photosynthesis to very diverse ecological niches and indicate a strong need for further and more detailed analyses. Such analyses should include e.g. phospho-proteomics of thylakoid membrane proteins of *Lemna* isolated from plants subjected to varying and also extreme illumination and nutrient conditions. We hypothesize that the basic evolutionary protein chassis of C3 photosynthesis in plant species with highly differing ecological niches remains well conserved at the sequence level, and that the required ecological adaptations to maintain photosynthesis efficiency as high as possible are achieved mostly by specific post-translational modifications. These are not restricted to phosphorylation alone and may contain also other types such as acetylation or glucosylation. More detailed studies of such modifications may provide important hints for improving indoor-farming approaches of terrestrial vegetables using aquaculture or for the plethora of biotechnology applications of *Lemnaceae*.

A side aspect of our study was the recording of high magnification videos of *Lemna minor* growth in a sugar-free liquid medium. The major obstacle for such video recordings is the floating of individual fronds on the medium (eventually moving out of the camera focus) making it difficult to record full time series of propagation in the long-term. We solved this problem by using plastic grids as anchor for the frond roots. While these grids largely immobilize the fronds they still allow a free-floating life style and normal vegetative budding. A recent study reporting a comparable imaging approach used cell culture plates with agarose-solidified medium as growth source and anchor for *Lemna minor* fronds [24]. While this approach resulted in effective immobilization of fronds and successful recordings of *Lemna minor* multiplication the resulting growth rates in sugar-free medium were relatively poor suggesting that solid growth media are not beneficial for *Lemna minor* growth. Growth rates determined in our system were largely indistinguishable from reported growth rates in liquid media. Our set-up thus provides a useful tool to perform high magnification live-imaging of *Lemnaceae* growth that is close to natural conditions and opens up a technological possibility to test the influence of ions, molecules or inhibitors on *Lemna minor* growth in high resolution.

## Materials and Methods

### Growth conditions and plant material

*Lemna minor* (strain 9441) was obtained from the Appenroth collection at the Friedrich-Schiller-University of Jena, Germany. Duckweed colonies were grown under long day conditions (8h dark/16h light) at 100 μmol photons m^-2^ s^-1^ white light provided by fluorescent stripe lamps (Lumilux Cool white L30W/840, Osram) at 21-24°C. Plant colonies were pre-cultured sterile in 1.5 l glass flasks sealed with cotton plugs (allowing for gas exchange) on nutrient media according to [13]. In order to maintain the cultures in the exponential growth phase, *Lemna* plants were regularly transferred to new flasks with fresh medium before they fully covered the available medium surface. For molecular and physiological analyses plant material was either harvested by pouring the media with the *Lemna* colonies through a sieve (for collection of larger biomass) or single colonies were transferred by hand with an inoculation loop into specified vessels (for details see results section) for *in vivo* Chl fluorescence measurements.

*Arabidopsis thaliana* wild-type (var. Columbia-0) was grown on potting soil under light and temperature conditions identical to those of the *Lemna* culture. For molecular analyses plant material of three to four-week-old rosettes was harvested directly under the growth lights, placed on ice or was flash-frozen in liquid nitrogen and immediately used for further analyses. For *in vivo* Chl fluorescence measurements plants were kept in their growth pots and placed under the emitter/detector unit (details see below) without any destructive manipulations.

### Growth rate determination by time lapse video imaging

*Lemna minor* fronds from the pre-culture were placed in a Petri dish equipped with a self-made, 3D-printer-generated grid and sealed with a gas-permeable tape under a digital microscope VHX 970F (Keyence, Japan) (compare Fig. 1C). Time lapse videos were recorded using the camera internal timer programme taking a picture every 15 min over the course of eight days resulting in 758 photos. Total video duration is 25 s with 30 frames per second speed. During video recording the fronds were maintained in a light/dark growth cycle comparable to the pre-culture by illuminating the Petri dish with two cold-white light sources (KL 1500 LCD, Zeiss, Germany) with an approximate intensity of 100-150 μmol photons m^-2^ s^-1^. Photos were taken during a 10 s illumination by the camera internal white LEDs. For determination of the growth rate of *Lemna minor* fronds the taken photos were digitally analysed for the leaf-covered area using freely available ImageJ software.

### *In vivo* chlorophyll fluorescence measurements

Room temperature chlorophyll fluorescence was detected by using a Junior pulse amplitude modulation (PAM) fluorometer (Walz, Effeltrich, Germany). For experimentation, individual fronds of *Lemna minor* were placed in a Petri dish containing growth media and immobilized by placing a weight on the tip of the roots to prevent plantlets from drifting on the liquid surface during the measurements. *Arabidopsis thaliana* (Col-0) plants grown on soil were measured for comparison. For both species, glass-fiber optics were placed 5 mm above the leaf surfaces in a way that the measuring light beam was centered on the leaf without touching the leaf edge. The intensity of the measuring light was adjusted to yield an approximate signal strength of 300 (rel. units) with non-dark-adapted plants. Plants were then dark-adapted for 30 minutes prior to each measurement. Actinic light intensity was set to 290 μmol *m^-2^ *s^-1^ photons of PAR, and quenching analysis was performed using the standard quenching and recovery protocol of the kinetics tab of the Junior-PAM software according to manufacturer’s protocols. Detected values were calculated within the software to provide the yield of various photosynthetic parameters (see results section).

For photo-inhibition experiments *Lemna minor* and *Arabidopsis thaliana* plants grown as described above were subjected to high light (HL) illumination treatments using a 300 W LED Grow Light panel for indicated time periods. The distance of the plants from the panel was adjusted to achieve a final PAR of approximately 1800 μmol m^-2^ s^-1^ photons at the leaf surface. For experimentation, individual fronds of *Lemna minor* were placed in 100 μl of growth media per well in black 96 multi-well plates (Greiner Bio-one) displaying a low auto-fluorescence thereby minimizing interference with the Chl fluorescence signal detection.

In order to obtain highest possible comparability between samples, individual leaves *of Arabidopsis* and individual wells with *Lemna fronds* were covered with light-tight aluminum foil before starting the high light treatment. The foil was removed successively at defined time points from the plants until the total exposure time (indicated in the figures) was reached. Chl fluorescence of individual leaves and plants was detected in parallel using a 2D Imaging-PAM fluorometer (Walz, Germany). To ensure maximal consistency between experiments and organisms, the multi-well plate containing *Lemna* fronds and *Arabidopsis* plant leaves were adjusted to the same distance from the camera lens. Before exposure to HL stress, plants were first dark-adapted for 30 minutes, and F_v_/F_m_ was measured as a reference. After high light treatment, plants were returned to their original growth light intensity and plants were allowed to recover from the light stress. F_v_/F_m_ was measured at three different time intervals (0h, 2h, and 19h) after HL treatment. To calculate the relative recovery, F_v_/F_m_ values obtained after HL treatment were expressed as a ratio to the reference values obtained before HL treatment. Each experiment was done in three independent biological and technical replicates (n=3).

### Western-immuno blot analyses

Freshly harvested material from *Lemna minor* and *Arabidopsis thaliana* was frozen in liquid nitrogen, homogenized in 1.5 ml Eppendorf tubes using pre-cooled pistils and mixed with equal volume of 4x SDS buffer (250 mM TRIS/ HCl pH 6.8; 40 % (w/v) glycerol; 8 % (w/v) SDS; 0.04 % bromophenol blue 5 % (v/v) 2-mercaptoethanol) per weight ground plant material. The samples were incubated for 5 min at 95°C. After centrifugation at 13,000 rpm for 10 min in an Eppendorf centrifuge, 5 μl of the supernatant containing the total soluble protein was separated by SDS–PAGE, using a 12 % gel. For initial loading adjustment the gel was stained with Coomassie and checked for sample comparability by judging the respective amount of lane loading. If required protein loading was adjusted by volume variation. Adjusted protein amounts were then transferred to a nitrocellulose membrane (Amersham™ Protran™ 0.2 μm NC, 10600001) by semi-dry blotting. After blocking the membrane with TRIS-buffered saline (TBS), containing 5% w/v skim milk powder overnight at 4°C, the membrane was washed twice with TBS-Tween (0.1 %) and once with TBS for around 10 min each. The membranes were then incubated for 2 h at room temperature with primary antibodies, diluted in TBS, containing 5 % milk powder [PSBA (Agrisera, AS05 084) 1:10,000; PSAA (Agrisera, AS06 172), FNR (Agrisera, AS15 2909) 1:5,000; PSBO (Agrisera, AS06 142-33) 1:5,000; PSBC (Agrisera, AS11 1787) 1:3000; LHCB1 (Agrisera, AS01 004) 1:2,000; ATPB (Agrisera, AS03 030) 1:5,000; PETA (Agrisera, AS20 4377) 1:1,000]. Membranes were washed three times for 10 min in TBS-Tween (0.1 %) and incubated for 1 h at room temperature with the secondary antibody, diluted in TBS, containing 5 % milk powder [goat anti rabbit or a Lumi IgG coupled to horse radish peroxidase (Agrisera, AS09 602) 1:25,000; or rabbit anti chicken IgY HRP (Promega Corporation, G1351) 1:10,000]. After washing (three times for 10 min in TBS-Tween (0.1 %) and twice for 10 min with TBS), the signal was detected using enhanced chemiluminescence (Lumi-Light^PLUS^ Western Blotting Substrate, Roche or Pierce®ECL2 Western Blotting Substrate) and a BIORAD camera system or a Lumi Imager F1 (Roche). Signal detection times were adjusted in a way that sufficiently strong signals above background were achieved. *Lemna* and *Arabidopsis* samples typically showed comparable reactivity to the antisera except the one for the D1 protein. Here the *Lemna* sample did react poorly requiring an extension of exposition time to reach the same signal strength as the *Arabidopsis* sample. This, however, had no consequences for the data interpretation.

### Protein sequence alignments

Alignments were performed, using BLASTP (NCBI). Sequences of *Arabidopsis thaliana* proteins (TAIR, UniProt) were used as a query. Standard parameter (standard databases, non-redundant protein sequences, threshold of 0.05, matrix BLOSUM62, cap costs existence: 11 and extension: 1 and conditional compositional score matrix adjustment) were used and it was blasted against *Lemna minor* (taxid:4472).

### 77K fluorescence emission spectra

77K low temperature fluorescence measurements to obtain Chl fluorescence of PSI relative to PSII were done in a custom-made device using an LED source and a CCD detection array (Ocean Optics) corresponding to described set-ups [25]. During the whole measurement, the sample containing part of the device was kept in the liquid nitrogen. All wavelengths between 650 nm and 850 nm were measured. First, a blank was measured to ensure, that the measuring cylinder was clean. Then one leaf was harvested, clamped into the measuring device and directly incubated in liquid nitrogen to ensure, that the leaf was completely frozen before the measurement was started. The measurement was repeated with the same leaf until two subsequent spectra resulted in the same values to ensure optimal measuring conditions. All recorded spectra were normalized to the average of the PSII Chl fluorescence emission peak at 686 nm. Three independent biological replicates for each condition were done and the standard deviation for the peaks at 686 and 735 nm were determined.

### Phosphorylation state of thylakoid proteins

For the analysis of the phosphorylation state, thylakoid membranes were isolated according to [26]. Samples were homogenized in 40 ml ice cold grinding buffer (50 mM HEPES/ KOH *pH* 7,5; 330 mM sorbitol; 2 mM EDTA; 1 mM MgCl_2_; 5 mM ascorbate; 0.05 % (w/v) BSA; 10 mM NaF), using a waring blender. The homogenate was filtered through filter tissue (50 μm pore size), centrifuged at 5,000 g and 4°C for 4 minutes (Sorvall, HB6) and the sediment gently solved in 20 ml of shock buffer (850 mM HEPES/ KOH *pH* 7,5; 5 mM sorbitol; 5 mM MgCl_2_; 10 mM NaF). After incubation for 4 min, samples were centrifuged at 5,000 g and 4°C for 4 minutes (Sorvall, HB6) and the sediment solved in 40 ml storage buffer (50 mM HEPES/ KOH *pH* 7.5; 100 mM sorbitol; 10 mM MgCl_2_; 10 mM NaF). Samples were further centrifuged at 5,000 g and 4°C for 4 minutes (Sorvall, HB6) and the sediment was solved in 750 μl or 1 ml storage buffer depending on the amount of fresh weight used. The chlorophyll concentration was measured according to [27]. Proteins were mixed with 4 x SDS sample buffer (0.075 M TRIS/ HCl *pH* 6.8; 6 M urea; 10% (v/v) SDS; 2 % (v/v) glycerol; 0.5 % (v/v) 2-mercaptoethanol; bromophenol blue) and denatured for 10 min at 68°C. Amounts corresponding to 1 μg chlorophyll were separated by SDS PAGE (12 %) and transferred to a nitrocellulose membrane (Whatman ®PROTAN Nitrocellulose Transfer Membrane 0.45 μm). The membrane was blocked for 1 h at room temperature with 5 % BSA dissolved in TBS-Tween (0.1 %) and incubated with the primary antibody rabbit anti-phospho-threonine [(Cell Signaling Technology®, #9381) 1:1000], diluted in 2 % (w/v) BSA in 1 x TBS-T (0.1 %) overnight at 4°C. The membrane was washed three times for 15 min in TBS-Tween (0.1 %), incubated with the secondary antibody rabbit anti chicken IgY HRP [(Promega Corporation, G1351) 1:10000] for 2 h and again washed three times. For detection, ECL solutions I (0.1 M TRIS/ HCl pH 8.5; 2.5 mM Luminol (in DMSO); 0.04 mM p-coumaric acid) and II (0.1 M TRIS/ HCl pH 8.5; 5.4 mM H_2_O_2_) were mixed 1:2 and applied for 2 min to the membrane. Signal detection was done for 7 sec or 1 min, respectively. For verification of equal protein loading, the membrane was applied to amido black staining after ECL detection. The membrane was incubated for 5 min in the staining solution (50 % (v/v) ethanol; 40 % (v/v) water II; 10 % (v/v) acetic acid; 4,06 mM Amido Black 10B) and de-stained in the de-staining solution (50 % (v/v) ethanol; 40 % (v/v) water II; 10 % (v/v) acetic acid).

## Supplementary materials

The growth series depicted in Fig. 1D represent static images at indicated time points. The full dynamic of *Lemna minor* growth becomes visible in the accompanying video. Video S1: Supplemental Video Lemna minor Growth. An additional text file contains all bioinformatics sequence analyses of the photosynthesis proteins tested by western-immuno-blotting (Fig. 3) including principle biochemical parameters, alignments and degrees of homologies. Text File S1: Supplemental Data File 1. All supplementary materials are available at https://doi.org/10.5281/zenodo.7817351.

## Acknowledgements

We are grateful to Klaus Appenroth who pointed our interest to duckweed research, provided the strains used in this study and gave us valuable advice in duckweed cultivation and physiology. Birgit Lippmann, Julia Gunia and Fabien Chevalier are acknowledged for their help in maintaining permanent duckweed cultures and providing technical support for molecular analyses. Giovanni Finazzi is acknowledged for his support in the 77K measurements. Maximilian Wiechmann and Thomas Gan are acknowledged for the construction of the 3D-printer-generated plastic grids.

## Author contribution

Conceptualization, M.L., S.O., T.P.; Methodology, M.L., S.O., T.P.; Formal Analysis, M.L., S.O., T.P.; Investigation, M.L., E.H., B.G., J.D., S.O., T.P.; Writing – Original Draft Preparation, M.L., S.O., T.P.; Writing – Review & Editing, M.L., S.O., T.P.; Supervision, M.L., S.O., T.P.; Project Administration, M.L., S.O., T.P.; Funding Acquisition, T.P.

